# Eye movements as a readout of sensorimotor decision processes

**DOI:** 10.1101/785832

**Authors:** Jolande Fooken, Miriam Spering

**Affiliations:** Dept. Ophthalmology & Visual Sciences, University of British Columbia, Vancouver, Canada; Graduate Program in Neuroscience, University of British Columbia, Vancouver, Canada; Center for Brain Health, University of British Columbia, Vancouver, Canada; Institute for Computing, Information and Cognitive Systems, University of British Columbia, Vancouver, Canada

**Keywords:** eye movements, prediction, perceptual decision making, visual motion, manual interception

## Abstract

Real-world tasks, such as avoiding obstacles, require a sequence of interdependent decisions to reach accurate motor outcomes. Yet, most studies on primate decision making involve simple one-step choices. Here we investigate how sensorimotor decisions develop over time. In a go/no-go interception task human observers (*n*=42) judged whether a briefly-presented moving target would pass (interception required) or miss (no hand movement required) a strike box while their eye and hand movements were recorded. Go/no-go decision formation had to occur within the first few hundred milliseconds to allow time-critical interception. We found that the earliest time point at which eye movements started to differentiate decision outcome (go vs. no-go) coincided with hand movement onset. Moreover, eye movements were related to different stages of decision making. Whereas higher eye velocity during smooth pursuit initiation (prior to “whether” decision) was related to higher go/no-go decision accuracy, faster pursuit maintenance was associated with accurate interception timing (“when” decision). These results indicate that pursuit initiation and maintenance are continuously linked to ongoing sensorimotor decision formation.

## New and Noteworthy

In this study we show that human eye movements are a continuous and sensitive indicator of go/no-go decision processes. We link different stages of decision formation to distinct oculomotor events during the open-loop vs. closed-loop phase of pursuit eye movements. Critically, the earliest time point at which eye movements started to differentiate decision outcomes coincided with hand movement onset, suggesting shared sensorimotor processing in the eye and hand movement systems. These results emphasize the potential of studying naturally occurring eye movements as a continuous read-out of cognitive processes.

## Eye movements as a readout of sensorimotor decision processes

Perceptual decisions in real-world scenarios often require a sequence of interdependent decisions. For example, when a pedestrian steps onto a bike lane, an approaching cyclist has to decide whether to stop or to veer around the obstacle. Depending on the initial decision outcome the cyclist then has to decide how hard to brake or in which direction to swerve. Dynamically-evolving decision processes have been studied in ecologically-inspired tasks, such as spatial navigation in rodents (Harvey et al., 2012; Krumin et al., 2018; Pfeiffer and Foster, 2013) or during visual search and foraging in human observers (Diamond et al., 2017; Najemnik and Geisler, 2005; Yoon et al., 2018). Yet, the time course of visually-guided sequential decisions in simple movement tasks is relatively unexplored. This study probes decision-making processes using a speeded manual go/no-go interception task. We investigate continuous eye movements during two-stage perceptual decisions as a key signature of the dynamics of decision making.

Goal-directed hand, arm, and body movements, such as those during obstacle avoidance, are accompanied by naturally occurring eye movements. During many natural tasks the eyes fixate on target objects as the hand approaches and shift to the next target at around the time the hand arrives (e.g., Ballard et al., 1992; Johansson et al., 2001; Land et al., 1999). Past research has consistently found a behavioural interdependency between eye and hand movements, indicating common or coordinated control of eye and hand motor control. Moreover, eye movements are sensitive indicators of sensorimotor decision formation and outcome (Fooken and Spering, 2019; Joo et al., 2016; McSorley and McCloy, 2009). For example, smooth pursuit and saccadic eye movement parameters provide reliable estimates of the outcome of go/no-go manual interception decisions in humans (Fooken and Spering, 2019). However, it is possible that eye movement modulations simply reflect observers’ actions rather than the perceptual decision. That is, the decision to act typically requires more accurate visual control than the decision not to act. Alternatively, eye movements may indicate an early readout of the decision formation itself, reflecting the observer’s response before it is executed. The current study examined the role of eye movements during the time course of a two-stage decision process: the decision whether and when to intercept a moving target.

## Material and Methods

### Overview

This paper relates eye movements to the time course of decision formation during a rapid go/no-go track-intercept task. To investigate the relationship between eye movements and task outcome we performed new analyses on a previously published data set (Fooken and Spering, 2019). Paradigm and procedure are identical to this published experiment and are reproduced here for the reader’s convenience. New analyses developed for the current paper are described in detail.

### Observers

We collected data from 45 male observers and excluded three participants who did not follow instructions and moved their hand in more than 80% of trials, regardless of stimulus conditions. The remaining 42 observers (mean age 20.5 ± 2.0 yrs) consisted of 25 members of the UBC male varsity baseball team and 17 age- and gender-matched non-athletes. All observers had normal or corrected-to-normal visual acuity confirmed by an ETDRS acuity chart; 39 were right-handed, six were left-handed (dominant hand was defined as the throwing hand). All observers were unaware of the purpose of the experiment. The experimental protocol adheres to the Declaration of Helsinki and was approved by the Behavioral Research Ethics Board at the University of British Columbia; observers gave written informed consent before participation.

### Visual display and apparatus

The visual target was shown at a luminance of 5.4 candela per meter squared (cd/m^2^) on a uniform grey background (35.9 cd/m^2^). Stimuli were back-projected onto a translucent screen with a PROPixx video projector (VPixx Technologies, Saint-Bruno, Canada; refresh rate 60 Hz, resolution 1280 (H) × 1024 (V) pixels). The displayed window was 44.5 (H) × 36 (V) cm or 55° × 45° in size. Stimulus display and data collection were controlled by a PC (NVIDIA GeForce GT 430 graphics card) and the experiment was programmed in Matlab 7.1 using Psychtoolbox 3.0.8 (Brainard, 1997; Kleiner et al., 2007; Pelli, 1997). Observers were seated in a dimly-lit room at 46 cm distance from the screen with their head supported by a combined chin- and forehead-rest.

### Experimental paradigm

Observers were asked to track a small moving target (diameter 2 deg) and to predict whether it would pass (“go” response required) or miss (“no-go” required) a designated strike box (**Fig. 1A,B**). We instructed observers to withhold a hand movement in miss trajectories and to intercept the ball while it was in the strike box in pass trajectories. Each interception started from a table-fixed position and was made with the index finger of the dominant hand.

**Figure 1.**
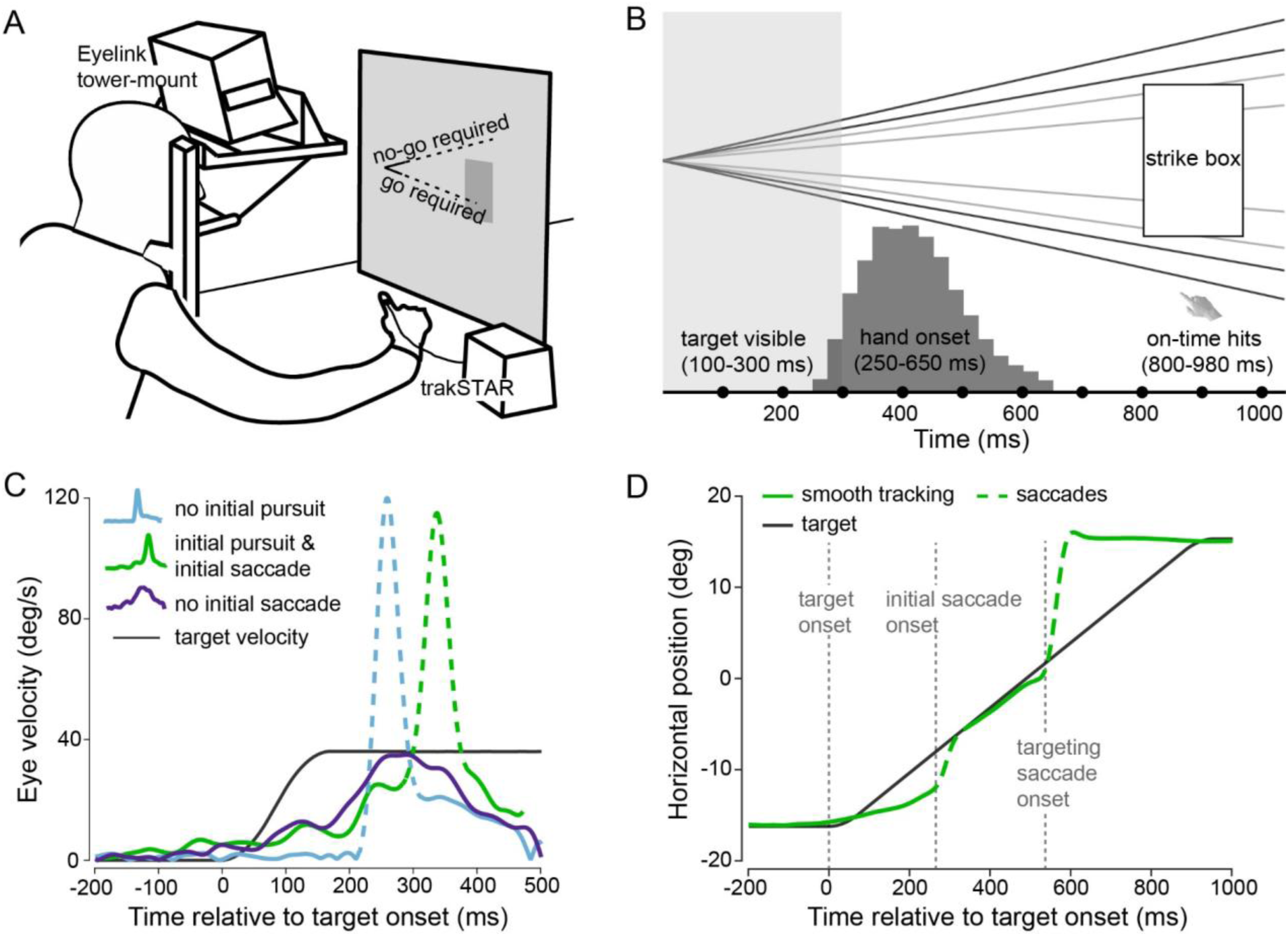
(A) Cartoon of the experimental setup. Observers had to judge whether a briefly presented target would pass (go required) or miss (no-go required) the strike box. Judgments were made by initiating or withholding an interceptive hand movement. (B) Task and interception events across time. The target was visible for 100-300 ms, entered the strike box at ∼800 ms, and was inside the box for ∼180 ms. Observers initiated hand movements between 250 and 650 ms. The trial ended when the target reached the end of its trajectory (no-go), or when observers intercepted it (go). (C) Exemplary initial eye velocity traces of single trials. Observers elicited three different types of eye movement patterns in response to target motion (black), either fixating until initiating a saccade (blue; 40.3% of trials), tracking the target before initiating a saccade (green; 53.9% of trials), or tracking it smoothly (purple; 5.8% of all trials). (D) Exemplary eye position trace of a single trial in which a combination of smooth pursuit (solid green line) and saccades (dashed green line) was exhibited. Target motion onset, initial saccade onset and targeting saccade onset are indicated by vertical dashed grey lines.

The stimulus followed a linear-diagonal path and either hit or missed a darker grey strike (31.5 cd/m^2^) box that was 6°×10° in size and offset by 12° from the center to the side of interception (**Fig. 1B**). Stimulus velocity followed natural forces (gravity, drag force, Magnus force; Fooken et al., 2016). Launch angles were set to ±5°, ±7° (pass trajectories), ±10°, or ±12° (miss trajectories). Target speed was either 36 or 41°/s. Importantly, the target disappeared shortly after launch (100-300 ms) making the task very challenging. All conditions were randomized and equally balanced. We instructed observers to track the target with their eyes and to follow its assumed trajectory even after it had disappeared. Each trial ended when observers either intercepted the target or when the target reached the end of the screen (1-1.1 s). At the end of each trial observers received feedback about their performance; target end position was shown, and correct or incorrect decisions were indicated. Each observer performed a familiarization session (16 trials; full trajectory visible) followed by 384 experimental trials in which the target viewing time was limited.

We defined four response types following conventions in the literature (Kim et al., 2005; Yang et al., 2010). Trials were classified as *correct go* if observers made an interception (i.e. touched the screen) in response to a pass trajectory and as *incorrect go* if observers moved their hands more than half way to the screen during a miss trajectory. Trials were classified as *correct no-go* or *incorrect no-go* if observers withheld a hand movement or moved their hand less than half way to the screen in response to a miss or pass trajectory, respectively. Decision accuracy was calculated as the percentage of all correct go and no-go responses.

### Eye and hand movement recordings and preprocessing

Eye position signals from the right eye were recorded with a video-based eye tracker (Eyelink 1000 tower mount; SR Research Ltd., Ottawa, ON, Canada) and sampled at 1000 Hz. Eye movements were analyzed off-line using custom-made routines in Matlab. Eye velocity profiles were filtered using a low-pass, second-order Butterworth filter with cutoff frequencies of 15 Hz (position) and 30 Hz (velocity). Saccades were detected based on a combined velocity and acceleration criterion: five consecutive frames had to exceed a fixed velocity criterion of 50°/s; saccade on- and offsets were then determined as acceleration minima and maxima, respectively, and saccades were excluded from pursuit analysis. Pursuit onset was detected in individual traces using a piecewise linear function that was fit to the filtered position trace (Fooken et al., 2016).

Finger position was recorded with a magnetic tracker (3D Guidance trakSTAR, Ascension Technology Corp., Shelburne, VT, USA) at a sampling rate of 240 Hz; a lightweight sensor was attached to the observer’s dominant hand’s index fingertip with a small Velcro strap. The 2D finger interception position was recorded in x- and y-screen-centered coordinates. Each trial was manually inspected and a total of 345 trials (2%) were excluded across all observers due to eye or hand tracker signal loss.

### Eye movement data analyses

The stimulus characteristics in this paradigm triggered tracking behavior that most closely resembled short periods of smooth pursuit and catch-up saccades (**Fig. 1C,D**). In some trials, observers tended to anticipate target motion (green and purple traces in **Fig. 1C)**, whereas in other trials, observers fixated until initiating a catch-up saccade to match target speed (blue trace in **Fig. 1C)**. To evaluate tracking behavior across time we analyzed eye movement quality during different time windows. In a previous study we found that observers typically made 2-3 saccades during the experimental paradigm. We defined the time interval from stimulus onset to the onset of the first saccade as our pursuit initiation time window and the time from first saccade offset to final saccade onset as the pursuit maintenance window (**Fig. 1D**). For trials, in which saccades occurred in between the first and last saccade we excluded the intermediate saccades from pursuit analysis. For the pursuit initiation and pursuit maintenance window we analyzed eye position and velocity relative to target position and velocity and extracted the following pursuit measures: mean eye and velocity position error, relative eye velocity (gain), and absolute eye velocity. For each observer, we analyzed mean saccade rate across time as a temporal measure that is independent of spatial target position. For each trial we created a vector aligned to stimulus onset that contained an assigned value of 1 (eye in saccade state) or 0 (eye in fixation or pursuit state) at each time point. The mean saccade rate was then determined by calculating the mean probability of the eye being in saccade state at each time point.

For the pursuit initiation interval we calculated a speed-accuracy score combining the latency of the initial saccade with initial pursuit velocity. We normalized eye velocity error *ê* (accuracy) and initial saccade latency 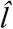 (speed) across all observers and trials. Note that we accounted for the inverse relationship between velocity error and accuracy (i.e. a higher velocity error corresponds to lower tracking accuracy) by calculating 1-*ê* as speed score. We then added the normalized speed and accuracy score and calculated an average speed-accuracy score

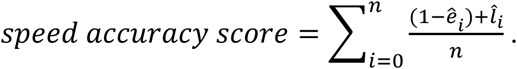

### Go/no-go separation time and statistical analyses

To calculate the time at which the eye movement signature starts to differ between go and no-go decision outcome, we calculated the saccade rate for each observer and split the data into go and no-go trials. We then compared the saccade rate between go and no-go decisions across time. We calculated a moving average of the saccade rate across a 5 ms time interval and down-sampled the data from 1000 Hz to 500 Hz to decrease the risk of detecting false negatives. We then performed a Mann-Whitney test for each time interval. The separation time was determined as the first time interval of at least three consecutive intervals for which a *p*-value smaller than 0.01 was achieved.

Differences between eye and hand movement measures were evaluated using Welch’s two-sample paired t tests. Correlations were assessed using the Pearson R test. All statistical analyses were performed in RStudio version 1.0.136 (RStudio, Boston, MA, USA).

## Results

Relating eye movements to decision formation and outcome (go/no-go) revealed three main findings. First, the earliest time point at which eye movements started to differentiate go/no-go decisions coincided with hand movement onset. This result indicates that differences in eye movements for go compared to no-go decisions were not merely a consequence of interceptive hand movements, but they occurred prior to hand movement execution. Second, higher eye velocity during pursuit initiation was related to higher decision accuracy (whether to intercept). Third, higher eye velocity during pursuit maintenance was related to better interception timing (when to intercept), suggesting that different stages of decision formation were linked to continuously evolving eye movements. In the following we will first qualitatively describe observers’ eye and hand movement response over the time course of the go/no-go interception task and then present quantitative results to support our three main findings.

### Eye movement separation coincides with hand movement onset

The go/no-go task employed in this study triggered a combination of smooth pursuit and saccadic eye movements. To determine at which time point eye movements differentiated go and no-go decisions (whether to intercept) we investigated the change in saccade rate—a temporal measure that is independent of the spatial target position—for go-compared to no-go decisions. Observers typically made 2-3 saccades in each trial (2.6 ± .4). The initial catch-up saccade (i.e. the first saccade in each trial) was on average elicited 240 ms (*SD* = 41.6 ms) after target onset followed by a brief period of tracking before a final, targeting saccade was made on average 620 ms after target onset (*SD* = 58.5 ms; **Fig. 1D**).

For each observer, saccade rates were compared between alternate decision outcomes (go vs. no-go). The time at which saccade rates first differed significantly (Mann-Whitney test, *p* < 0.01) was determined to be the go/no-go separation time (see Materials and Methods; **Fig. 2A and Figure supplement 1A,B**). Using this method, we were unable to find a separation time for three observers until after the offset of the final saccade (**Figure supplement 1B,C**). Separation times for these observers differed by two or more standard deviations from the group mean and were excluded from this part of the analysis. For the remaining 39 observers the mean separation time was 395 ms (range 326-520 ms; **Fig. 2B**). In go trials, the same observers initiated a hand movement on average around the same time at 411 ms (range 320-536 ms; separation time vs. hand movement onset: *t*(38) = 2.4, *p* = .02; **Fig. 1B**). These results indicate that the time at which eye movements start to reflect decision outcome coincides with hand movement onset. Therefore, eye movement patterns that reflect go/no-go decisions are not simply a consequence of hand movement execution, but an indicator of the ongoing decision formation.

**Figure 2.**
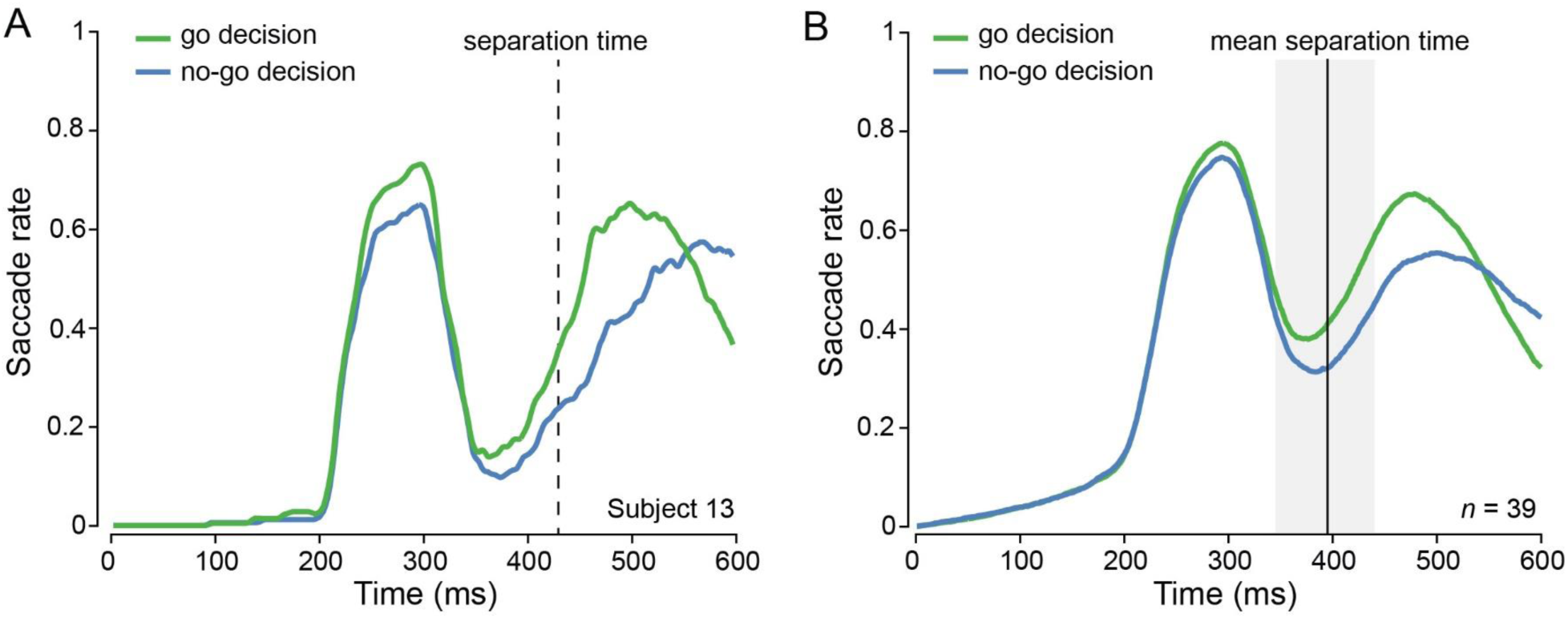
Eye movement separation for go vs. no-go decisions. (A) Saccade rates for go vs. no-go decisions of a representative observer with a separation time of 428 ms. (B) Saccade rates for go vs. no-go decisions averaged across all observers that showed a differentiation (*n* = 39).

### Pursuit initiation is related to decision accuracy

The time course of eye and hand movements suggests that go/no-go decision formation occurred within the first few hundred milliseconds of each trial. The initial saccade offset was on average 350 ms (*SD* = 41.1 ms) after stimulus onset, only 50-ms before the average onset of interceptive hand movements. The brief delay between initial saccade offset and hand movement onset suggests that interception decision formation occurred prior to the initial saccade. We next investigated whether pursuit initiation (target onset to initial saccade onset) was related to go/no-go decision accuracy. Faster eye movements during the pursuit initiation period were associated with more accurate decisions (**Fig. 3A**), reflected in a significant positive correlation between pursuit initiation velocity and decision accuracy (*r* = .51, *p* < .001). Congruently, average eye velocity error (2D velocity difference between eye and target) during pursuit initiation was negatively correlated with decision accuracy (*r* = -.39, *p* = .01; **Fig. 3B**). These results suggest that the initiation of smooth pursuit and the resulting decrease in velocity error might be related to target motion prediction and decision formation accuracy.

**Figure 3.**
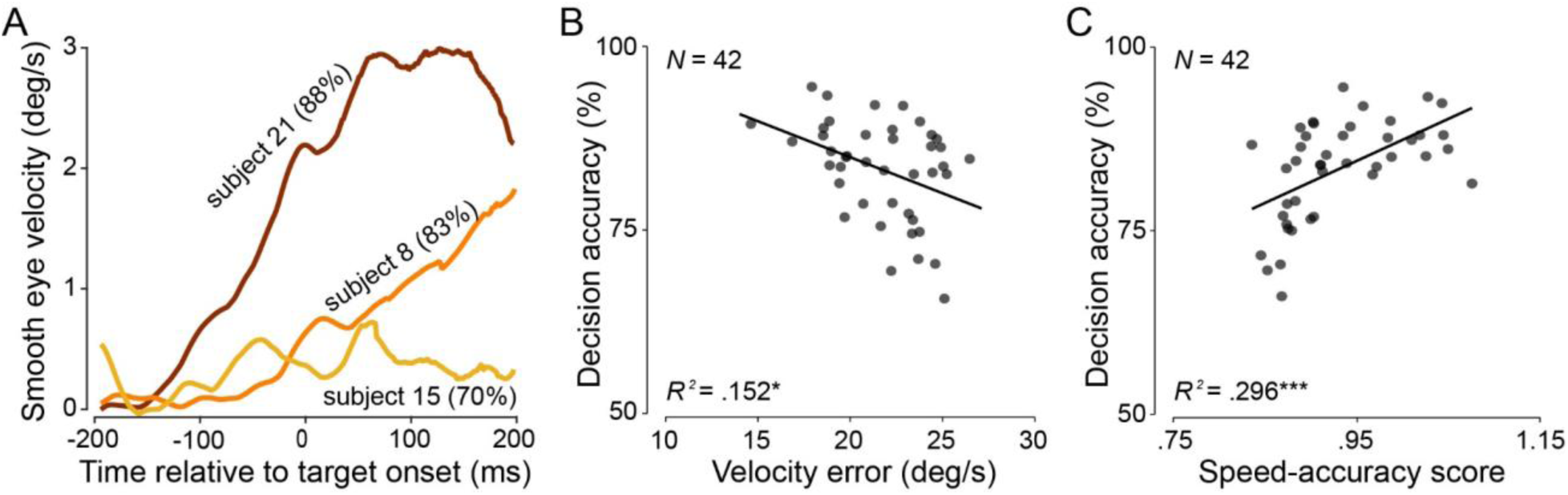
Relationship between eye movement initiation and decision accuracy. (A) Initial eye velocity from three observers averaged across 384 trials. Subject 15 (yellow) shows the lowest eye velocity during the pursuit initiation phase and had an overall lower decision accuracy (70%) than observers 8 (orange; 83%) and 21 (brown; 88%). (B) Decision accuracy is negatively related to eye velocity error in the interval from target onset to initial saccade onset. Asterisks denote significant regression results; * *p* < .05, *** *p* < .001. (C) Decision accuracy is positively related to modelled speed-accuracy score. Each data point represents the averaged value per observer.

However, we found that in ∼40% of all trials observers fixated until initiating the first catch-up saccade (**Fig. 1C**). In these trials, observers might benefit from delaying the initial catch-up saccade to allow more time for evidence accumulation, resulting in a potential speed-accuracy trade-off. To investigate whether initial saccade timing can account for some of the variability observed in the relationship between velocity error and decision accuracy (**Fig. 3B**), we calculated an average speed-accuracy score for each participant (see Material and Methods), and related this score to each observer’s decision accuracy. The observed positive correlation between the speed-accuracy score and decision accuracy (*r* = .54, *p* < .001; **Fig. 3C**) indicates that the timing of the initial saccade plays an important role in decision formation accuracy.

To further investigate the role of the accuracy and timing of pursuit initiation within subjects we divided observers into two groups—one that appeared to rely on reducing velocity error (group 1) and one that seemed to delay the initial saccade (group 2). Five observers did not reliably initiate smooth pursuit (<10% of the trials) and were automatically assigned into the saccade delay group (group 2). For the remaining observers we performed a median split analysis on initial eye velocity error and initial saccade and assigned observers to the group for which the change of decision accuracy between lower and upper bound was greater. We repeated the same analysis but instead of calculating a median split per observer used set cut-off values (23 deg/s; 230 ms) based on the entire sample. Five observers were classified into the respective other group, but overall patterns remained unchanged. For group 1, we found that decision accuracy was significantly higher in trials with a low compared to a high velocity error (*t*(23) = 4.1, *p* < .001; **Fig. 4A**). For group 2 we found that decision accuracy was higher for late as compared to early initial saccade latencies (*t*(17) = 2.7, *p* = .01; **Fig. 4B**). These results suggest that the timing and accuracy of pursuit initiation is related to go/no-go decision accuracy across as well as within observers.

**Figure 4.**
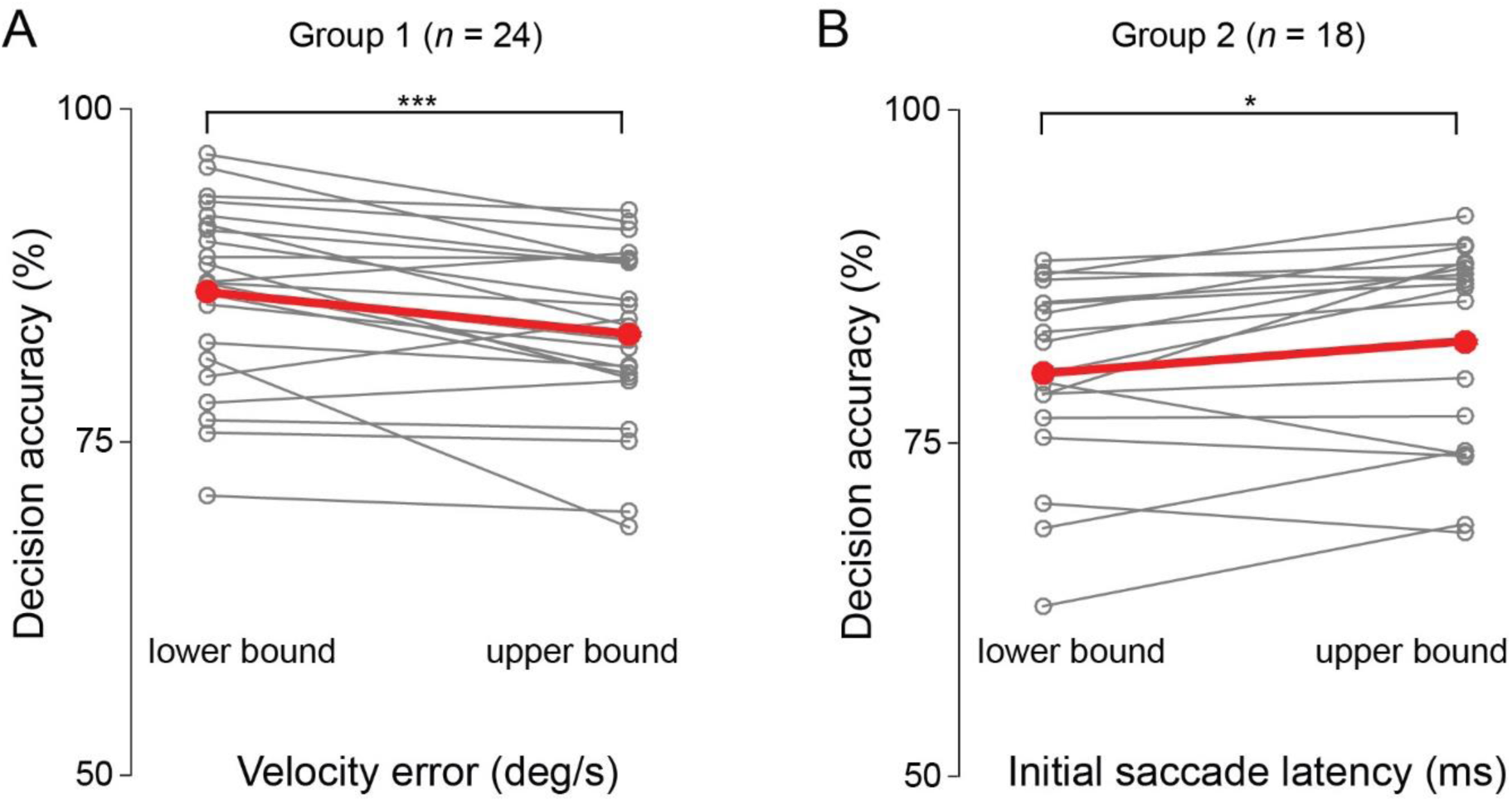
Comparison of go/no-go decision accuracy within each subject. (A) Difference in decision accuracy between trials in which eye velocity error was lower or higher than the median value for each observer (*n* = 24). (B) Difference in decision accuracy between trials in which initial saccade latency was later or earlier than the median value for each observer (n = 18). Grey lines show individual subject data, red thick line indicates group mean.

### Pursuit maintenance is related to hitting accuracy

The previous results indicate that eye movement initiation is linked to the first stage of go/no-go decisions (whether to intercept). In our task, go-decisions were always associated with a second decision: when to intercept. Observers were instructed to hit the strike box while the target was inside. Whereas the spatial position of the target trajectory was restricted to the area of the strike box, observers had to time-critically judge horizontal target motion to successfully intercept the target. We therefore focused on interception timing as a measure of interception accuracy (**Fig. 5A**). Across all interception (go-required) trials, observers’ interceptions were on time in 76 ± 6.6% of trials, too early in 18 ± 7.5% trials, and too late in 6 ± 4.5% trials. Incorrectly timed— early and late—interceptions were reflected in shifts in interceptive hand and eye movement onsets (**Fig. 5B,C**; **Table 1**). In trials in which they intercepted too early as compared to trials in which they were on time, observers moved their hand earlier (*t*(41) = 15.3, *p* < .001) and faster (*t*(41) = 4.9, *p* < .001) and initiated the final targeting saccade earlier (*t*(41) = 6.3, *p* < .001). Conversely, when interceptions were too late vs. on time, observers initiated the interceptive hand movement later (*t*(41) = 11.9, *p* < .001) and made the final targeting saccade later (*t*(41) = 4.1, *p* < .001), whereas the finger velocity did not differ (*t*(41) = 0.1, *p* = .89).

**Table 1:** Hand and eye movement differences for early, on-time, and late hits. Hand and targeting saccade latencies are relative to stimulus onset. The targeting saccade was defined as the final saccade of each trial.

|  | Hand latency | Hand peak velocity | Targeting sac. latency |
| --- | --- | --- | --- |
| Early hit | $372.4 \pm 38.2$ ms | $59.7 \pm 6.2$ cm/s | $610.1 \pm 57.5$ ms |
| On-time | $426.8 \pm 44.4$ ms | $57.3 \pm 5.5$ cm/s | $642.3 \pm 55.1$ ms |
| Late hit | $513.1 \pm 60.8$ ms | $57.4 \pm 6.9$ cm/s | $678.9 \pm 80.6$ ms |

Values indicate group averages ± standard deviation.

**Figure 5.**
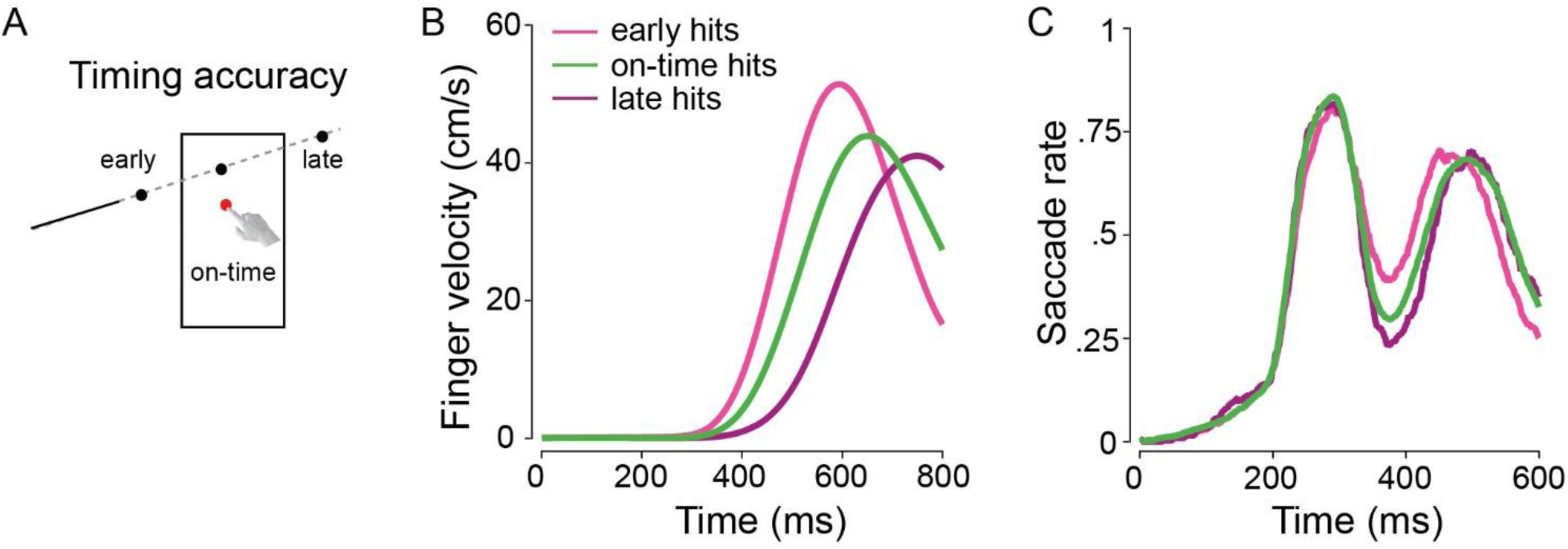
(A) Timing accuracy depended on the timing of the interception. If the target had not yet reached the strike box at the time of interception the observer was too early; if it had already left the strike at the time of interception the observer was too late. Finger velocity (B) and saccade rate (C) separated by early (light pink), on-time (green), and late (dark pink) hits reflect timing errors.

**Figure 6.**
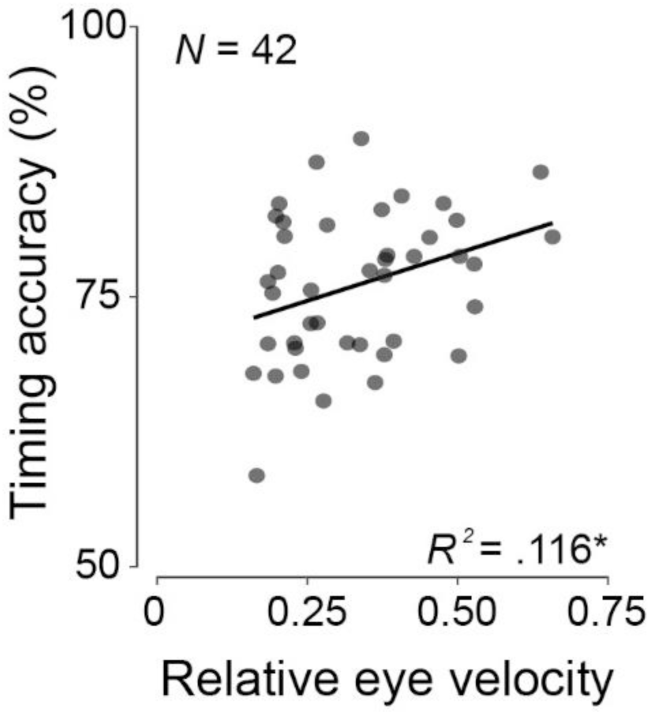
Relationship between timing accuracy and relative eye velocity during maintenance phase.

To investigate the relationship between eye movements and interception timing accuracy we analyzed eye movement velocity relative to target velocity during the pursuit maintenance phase (initial saccade offset to targeting saccade onset; **Fig. 1C**). We observed a positive correlation between relative eye velocity and interception timing accuracy (*r* = .34, *p* = .03; **Fig. 5B**). Positional measures of tracking quality (2D or horizontal eye position error, saccade amplitudes) were not related to timing accuracy. These results indicate that observers benefit from matching eye and target velocity during pursuit maintenance when tasked to accurately judge target speed and successfully time an interception.

## Discussion

In this study we related continuously evolving eye movements to two-stage perceptual decisions in a go/no-go interception task. We showed that eye movements distinguished go/no-go decisions early in the decision process and were not merely a consequence of motor execution. We also revealed that accurate smooth pursuit initiation was related to decision accuracy and that accurate smooth pursuit maintenance was linked to interception timing accuracy. These findings suggest that smooth pursuit eye movements continuously contribute to dynamic decision formation.

### Eye movements as an early indicator of go/no-go decisions

Eye movements are closely related to cognitive goals in a variety of everyday tasks that require an interaction with objects, such as brick stacking or sandwich making (Hayhoe, 2017; Hayhoe and Ballard, 2005; Land et al., 1999). A particularly strong link between eye movements and action is seen in the context of goal-directed hand movements. It is commonly observed that the eye leads the hand when tasks require pointing, hitting, or catching (Bekkering et al., 1994; Belardinelli et al., 2016; Brenner and Smeets, 2011; Land and McLeod, 2000; Mrotek and Soechting, 2007). We recently showed that eye movements reliably decoded go/no decisions; go compared to no-go decisions were associated with earlier targeting saccades to guide the interceptive hand movement (Fooken and Spering, 2019).

The current study goes beyond previous work by addressing the question whether eye movements are simply the consequence of a perceptual decision or if they reflect decision formation over time. Our results reveal that eye movements differentiate between later decisions whether or not to intercept at an early point in time, simultaneous with the onset of the interceptive hand movement. This finding emphasizes that eye movements indicate go/no-go decisions before hand movements are executed. The concurrence of eye movement separation time and hand movement onset is further evidence for common neural processing of action goals (e.g., Andersen and Cui, 2009; Crawford et al., 2004; Hwang et al., 2014). Moreover, our findings are closely related to the observation that eye and hand movements are interdependent during movement planning (Leclercq et al., 2013) and execution (Danion and Flanagan, 2018; Fooken et al.; 2016, 2018).

### Pursuit eye movements are related to decision accuracy and timing

We show that eye velocity during pursuit maintenance is linked to accurate interception timing. Previous research has shown that engaging in smooth pursuit aids accurate motion prediction, a benefit that is thought to arise from additional motion information provided through efference copy signals during pursuit maintenance (Bennett et al, 2010; Spering et al., 2011). Moreover, observers’ speed perception critically depends on the rate and direction of corrective saccades during tracking. Compared to trials in which observers tracked the target with pure smooth pursuit, observers overestimated target speed when tracking was accompanied by forward saccades, and underestimated target speed when backward saccades were elicited (Goettker et al., 2018). Corrective saccades during smooth pursuit also affected manual interception accuracy: observers intercepted ahead or behind of the target when eliciting forward or backward saccades, respectively (Goettker et al, 2019). In our task, target motion was predictable and only forward saccades were elicited. We did not find any relationship between saccade rate or amplitude and accurate interception timing. Instead, we found that relative eye velocity with respect to the target velocity, was linked to timing accuracy. These results complement previous findings showing that more accurate smooth pursuit eye movements (lower 2D position error) were linked to spatially more accurate manual interceptions (Fooken et al., 2016). Taken together these results suggest that velocity and positional error signals during smooth pursuit eye movements may contribute to different aspects of motion perception.

We further found that both saccades and smooth pursuit measures were related to decision timing and accuracy. These results highlight the interdependence of the two eye movement systems and indicate that smooth pursuit and saccades work in synchrony to enable accurate motion prediction (Barborica and Ferrera, 2004; Blohm et al, 2003; de Brouwer et al., 2002; Orban de Xivry et al., 2006; Orban de Xivry and Lefèvre, 2007; Schreiber et al., 2005). Our findings show that an increase in eye velocity as well as timing of the initial saccade are both beneficial for go/no-go decision accuracy, a novel finding that needs to be investigated further to identify the underlying mechanisms and speed-accuracy trade-offs.

### Perceptual decision making and hand motor responses are interdependent

Perceptual decisions can be biased by motor actions. For example, when participants indicated their choice in a motion discrimination task by left- or right-handed reaches that were associated with different mechanical loads, their motion perception was biased towards the side that had lower resistance. Interestingly, this perceptual bias occurred even though participants were not aware of the difference in motor cost between the two hands (Hagura et al., 2017). Notwithstanding these biases, eye and hand movements are modulated by prior perceptual decisions. When observers made visually guided (Joo et al., 2016) or choice-indicating (McSorley and McCloy, 2009) saccades just after a perceptual judgement, saccades in the decision-congruent direction were initiated earlier and faster. When observers’ hand movements were perturbed while making manual choice responses in a motion discrimination task, arm muscular reflex gains scaled with stimulus motion strength (Selen et al., 2012). This finding suggests that sensorimotor control is linked to ongoing perceptual decision making. Taken together, these findings indicate that there is a continuous crosstalk between perceptual decision processes and evolving motor plans.

Further evidence for the close relationship between perceptual and motor processing during decision making comes from studies of neural activity in motor cortex in human and non-human primates. Neural population activity measured by magnetoencephalography in human observers were predictive of decision outcome in a motion detection task before observers indicated their choice (Donner et al., 2009; Pape and Siegel, 2016). Furthermore, electrophysiological recordings of the dorsal premotor and primary motor cortex of macaque monkeys revealed that neural activity reflects changes of mind during reach target selection when the position of correct targets had to be updated dynamically (Kaufman et al., 2015; Thura and Cisek, 2014). These results suggest that the readout of sensory information is continuously coupled to motor preparation and execution.

### Cortical decision correlates

Neural and behavioural correlates of perceptual decision making have classically been studied using random-dot motion stimuli gradually adding to our understanding of decision networks in human observers (Gold and Shadlen, 2007; Heekeren et al., 2008; Schall, 2013). Yet, real world scenarios require more complex perceptual decisions than judging net motion. In a sequential decision task non-human primates were trained to select a target that was associated with a certain rule (pick the smaller or darker target). Monkeys then had to discriminate two visual targets based on their initial choice and responded by making a saccade to the chosen target (Abzug and Sommer, 2018). Rule selection and sequential decision monitoring were related to neural activity in the supplementary eye field, an area also associated with the predictive control of eye movements (Fukushima et al., 2006).

Similarly, a series of seminal studies investigating go/no-go decisions in human (Heinen et al., 2006) and non-human primates (Kim et al., 2005; Yang and Heinen, 2014; Yang et al., 2010) revealed neural decision correlates in the supplementary and frontal eye fields. Speed-accuracy trade-off of saccadic eye movements in a visual search task is also encoded in the frontal eye fields (Heitz and Schall, 2012). Taken together, these findings indicate that neural activity in the supplementary and frontal eye fields governs timing and performance monitoring of visual decision making and may play a key role in our paradigm.

### Limitations

One limitation of using a go/no-go paradigm is that decision accuracy is a binary variable, that is, the decision to go (or not to go) is either correct or incorrect. Designing a task with a continuous measure of decision accuracy would allow us to carry out a more detailed trial-by-trial analysis than the median split analysis presented here (**Fig. 5**). Yet, go/no-go decisions are interesting to study because the motor response is all-or-none and decisions cannot be corrected online. Another consideration is that manual interceptions had to occur within a specific time window. Hand movement onset or interception time can therefore not be interpreted as a classic measure of reaction time. The effect of decision timing on hand movement reaction time could be investigated in a future study. Notwithstanding these limitations, our results provide evidence for an interdependency of eye and hand movements with sensorimotor decision processes in human observers.

### Conclusion

Our findings emphasize commonalities in the timing and accuracy of oculomotor and hand movement control during decision-making. Eye movements provide a continuous readout of cognitive processes during two-stage decision formation. Because eye movements occur naturally and spontaneously, this may open new avenues for studying decision making in real-world scenarios.

## Acknowledgements

This work was supported by NSERC Discovery and Accelerator Grants (RGPIN 418493) and a Canada Foundation for Innovation John R. Evans Leaders Fund equipment grant to MS. The authors thank Rose Shannon for help with data collection and preprocessing, and Terry McCaig and the UBC varsity baseball team for their collaboration. Data were presented in preliminary form at the 2019 Vision Sciences Society meeting in St Pete’s Beach, FL (Fooken and Spering, Program No.: 21.12) and the 2019 Gordon Research Seminar on Eye Movements in Lewiston, ME.

## Supplementary material

**Figure supplement 1.**
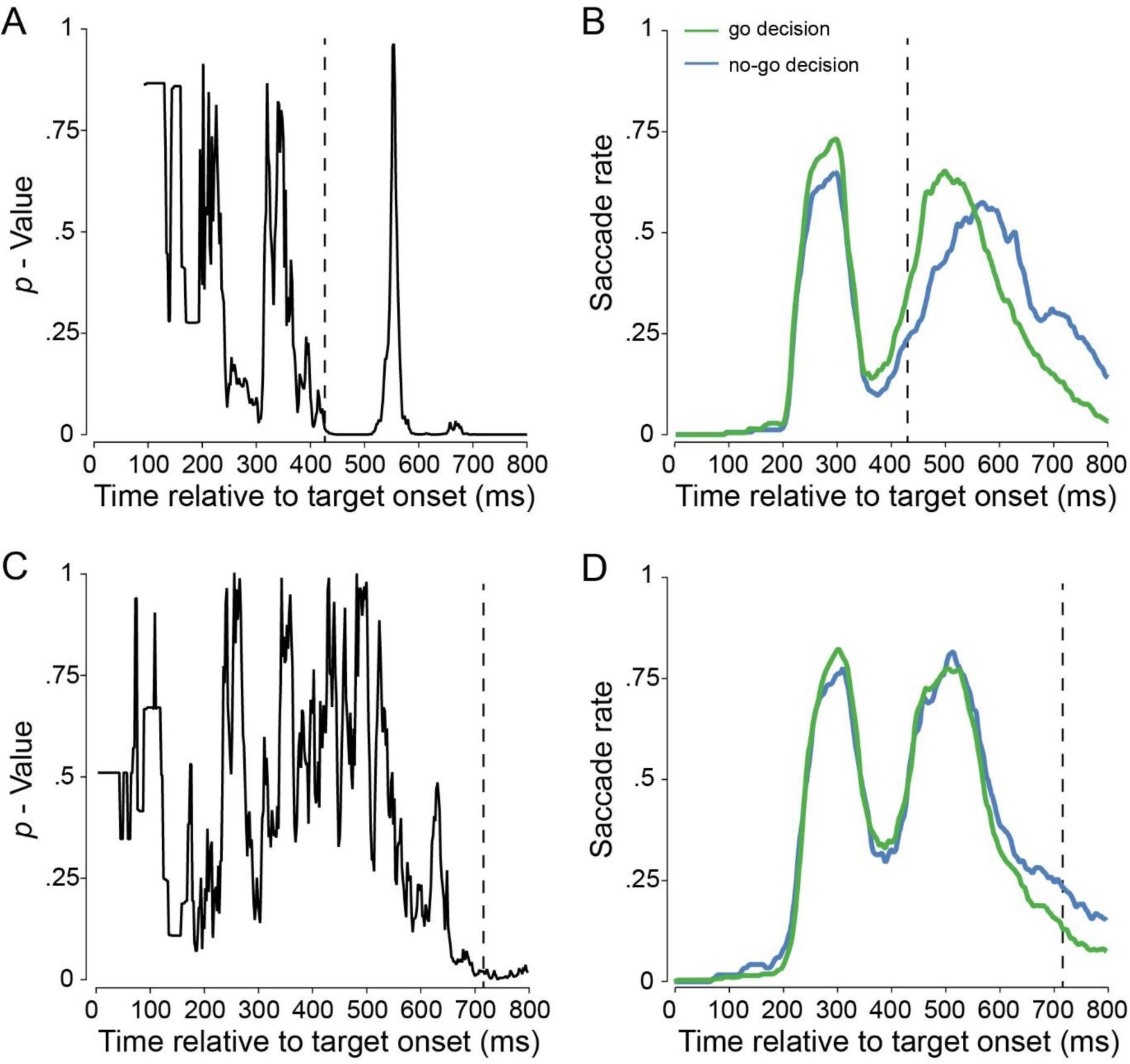
Calculation of separation time for two individual observers. (A) Change of *p*-value over time. First time point at which *p* < 0.01 is indicated by vertical dashed line. (B) Corresponding change of saccade rate over time. Eye separation time for this observer was at 428 ms. (C) Change of *p*-value and (D) saccade rate for an observer that did not show differential eye movements for go compared to no-go decisions.

